# Identification of functional Spo0A residues critical for sporulation in *Clostridioides difficile*

**DOI:** 10.1101/2022.02.07.479450

**Authors:** Michael A. DiCandia, Adrianne N. Edwards, Joshua B. Jones, Grace L. Swaim, Brooke D. Mills, Shonna M. McBride

## Abstract

*Clostridioides difficile* is an anaerobic, Gram-positive pathogen that is responsible for *C. difficile* infection (CDI). To survive in the environment and spread to new hosts, *C. difficile* must form metabolically-dormant spores. The formation of spores requires activation of the transcription factor Spo0A, which is the master regulator of sporulation in all endospore-forming bacteria. Though the sporulation initiation pathway has been delineated in the Bacilli, including the model spore-former *Bacillus subtilis*, the direct regulators of Spo0A in *C. difficile* remain undefined. *C. difficile* Spo0A shares highly conserved protein interaction regions with the *B. subtilis* sporulation proteins Spo0F and Spo0A, although many of the interacting factors present in *B. subtilis* are not encoded in *C. difficile*. To determine if comparable Spo0A residues are important for *C. difficile* sporulation initiation, site-directed mutagenesis was performed at conserved receiver domain residues and the effects on sporulation were examined. Mutation of residues important for homodimerization and interaction with both positive and negative regulators of *B. subtilis* Spo0A and Spo0F impacted *C. difficile* Spo0A function. The data also demonstrated that mutation of many additional conserved residues altered *C. difficile* Spo0A activity, even when the corresponding *Bacillus* interacting proteins are not apparent in the *C. difficile* genome. Finally, the conserved aspartate residue at position 56 of *C. difficile* Spo0A was determined to be the phosphorylation site that is necessary for Spo0A activation. The finding that Spo0A interacting motifs maintain functionality suggests that *C. difficile* Spo0A interacts with yet unidentified proteins that regulate its activity and control spore formation.

## INTRODUCTION

Sporulation initiation is a complex developmental process that allows for prolonged survival when environmental conditions become unfavorable. Some members of the Firmicutes phylum transition into metabolically dormant endospores (spores) that remain inert until environmental conditions are favorable again and the spore germinates to produce vegetative cells. Sporulation is energetically costly and, as such, highly regulated [1-5]. Sporulation initiation is controlled by the conserved transcription factor, Spo0A, the essential regulator of the sporulation gene expression program. Spo0A is encoded in all endospore-forming species and is regulated by phosphorylation of a conserved aspartate residue [6]. In the activated form, phosphorylated Spo0A undergoes a conformational change that facilitates self-dimerization. Activated Spo0A can then bind specific promoter regions, referred to as “0A boxes”, to regulate gene expression and trigger entry into the sporulation pathway [7, 8].

Sporulation initiation has been extensively studied in the model spore-former, *Bacillus subtilis*. In *B. subtilis* and other Bacilli, the phosphorylation status of Spo0A is controlled through a multicomponent phosphorelay, with the orphan sensor histidine kinases, KinA, KinB, KinC, KinD, and KinE, transferring phosphate to the intermediate phosphotransferase, Spo0F. Spo0F in turn mediates the flow of phosphate to the phosphotransferase Spo0B [9]. Spo0B then directly phosphorylates Spo0A, which activates sporulation-specific gene expression [2]. The Rap phosphatases, such as RapA, RapB, and RapH, can dephosphorylate Spo0F, while the Spo0E family of proteins dephosphorylate Spo0A, adding additional control to the phosphorelay (**Figure 1A**). The ability of Spo0B to interact with both Spo0F and Spo0A at shared, highly conserved motifs suggests a critical role for these residues in the regulation of Spo0A activity [10, 11].

**Figure 1.**
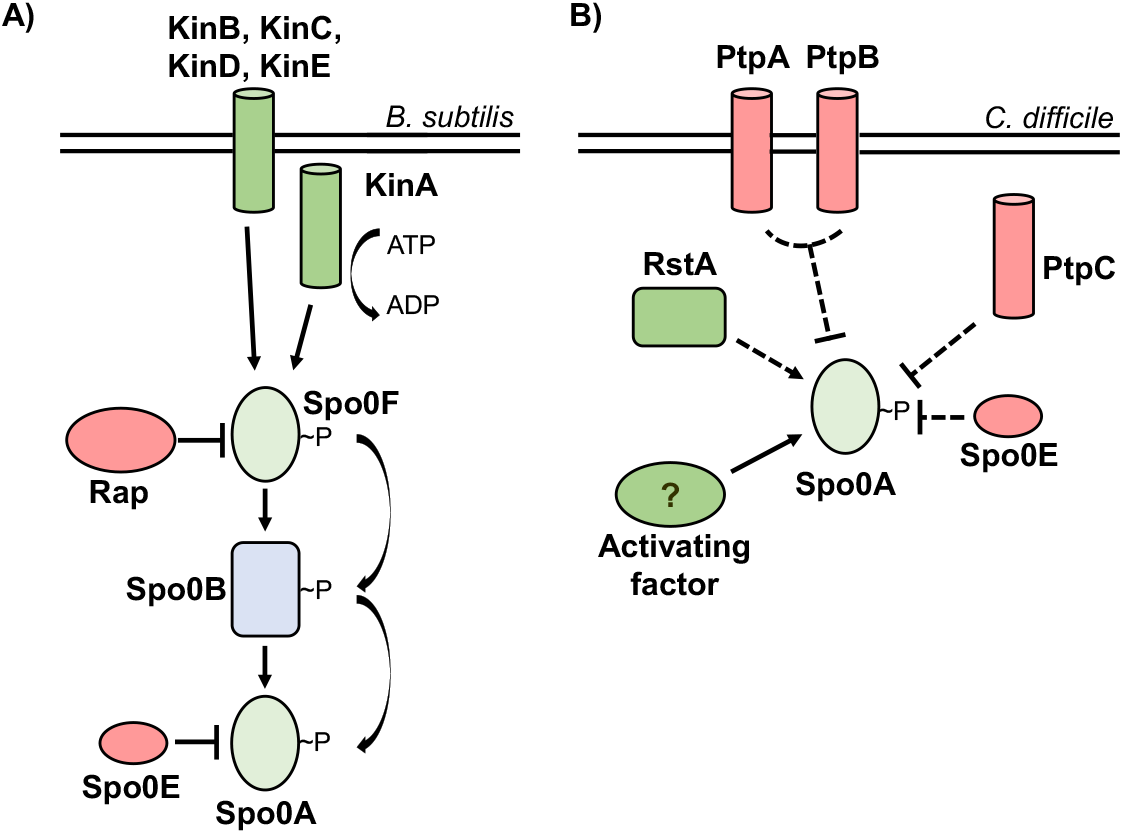
Evolutionarily divergent strategies for Spo0A activation. **A)** In *Bacillus* species, Spo0A is activated via the phosphorelay, with kinases KinA, KinB, KinC, KinD, and KinE transferring phosphate to Spo0A via Spo0F and Spo0B. The Rap and Spo0E phosphatases repress Spo0A activation through dephosphorylation of Spo0F and Spo0A, respectively. **B)** In *C. difficile*, the phosphotransfer proteins PtpA and PtpB act in coordination to prevent Spo0A activation. PtpC and Spo0E also repress Spo0A activity. RstA promotes sporulation through an unknown mechanism, and an unidentified activating factor is hypothesized to phosphorylate Spo0A.

Like the Bacilli, all spore-forming members of the anaerobic Clostridia encode *spo0A* [12]. However, the mechanisms of Spo0A regulation in the Clostridia, including *C. difficile*, are poorly characterized. The Spo0F-Spo0B phosphorelay is not apparent in clostridial genomes, suggesting that there are divergent mechanisms of Spo0A activation [13]. In some clostridial species, phosphotransfer proteins interact directly with Spo0A to activate or inactivate sporulation in a manner consistent with a traditional two-component system [14-17]. *C. difficile* encodes five orphan putative histidine kinases, three of which resemble the *B. subtilis* Spo0A-associated kinases and negatively regulate sporulation (PtpA, PtpB, and PtpC), and two that are not involved in sporulation [18, 19]. While one orphan kinase, PtpC, was reported to phosphorylate Spo0A *in vitro* [20], it was recently shown that a *ptpC* null mutant exhibits variably increased sporulation, demonstrating that PtpC negatively impacts Spo0A activity in the conditions tested [18] (**Figure 1B**). As none of the *C. difficile* orphan kinases are verified activators of Spo0A, it is challenging to predict the specific strategy of *C. difficile* Spo0A regulation.

Although Spo0F and Spo0B are not found in *C. difficile*, the regions of the Spo0A receiver domain that interact with these and other *Bacillus* regulators appear to be conserved in *C. difficile*. We hypothesized that conserved *Bacillus* Spo0A and Spo0F residues are also functionally important for *C. difficile* Spo0A regulation. To better understand how *C. difficile* Spo0A activity is regulated, we performed site-directed mutagenesis of conserved regions of the receiver domain that are functionally important for *B. subtilis* Spo0A and Spo0F and examined the effects on sporulation. Here we report on the residues and potential interaction surfaces that are important for regulation of *C. difficile* Spo0A activity.

## RESULTS

### The Spo0A *B. subtilis* and *C. difficile* N-terminal receiver domains are highly conserved

In *B. subtilis*, Spo0A and Spo0F share similar response regulator receiver domains (residues 6 – 116 in Spo0F and 6 – 120 in Spo0A), and both proteins interact with the phosphotrasfer protein Spo0B using conserved secondary structure [10, 21-24]. The residues of *B. subtilis* Spo0F and Spo0A that are important for signal transduction were previously identified and characterized [4, 10, 11, 25-28]. We aligned the amino acid sequence of the *B. subtilis* Spo0A (**Figure 2**) and Spo0F receiver domains (**Supplementary Figure 1**) to *C. difficile* Spo0A to predict orthologous functional residues. After identifying corresponding functional residues in the *C. difficile* Spo0A amino acid sequence, we performed site-directed mutagenesis of 30 *C. difficile* Spo0A residues to alanine, with the exception of native alanine residues, which were mutated to serine (**Figure 2B**). Mutated *spo0A* alleles driven by the *C. difficile spo0A* native promoter were expressed in a *C. difficile spo0A* mutant [29]. To assess the stability of mutant Spo0A proteins, we performed western blotting using an anti-Spo0A antibody [30] and found that all mutant Spo0A proteins, except those containing the Q17A, V18A, and P60A mutations, were stable under sporulating conditions (**Supplementary Figure 2**).

**Figure 2.**
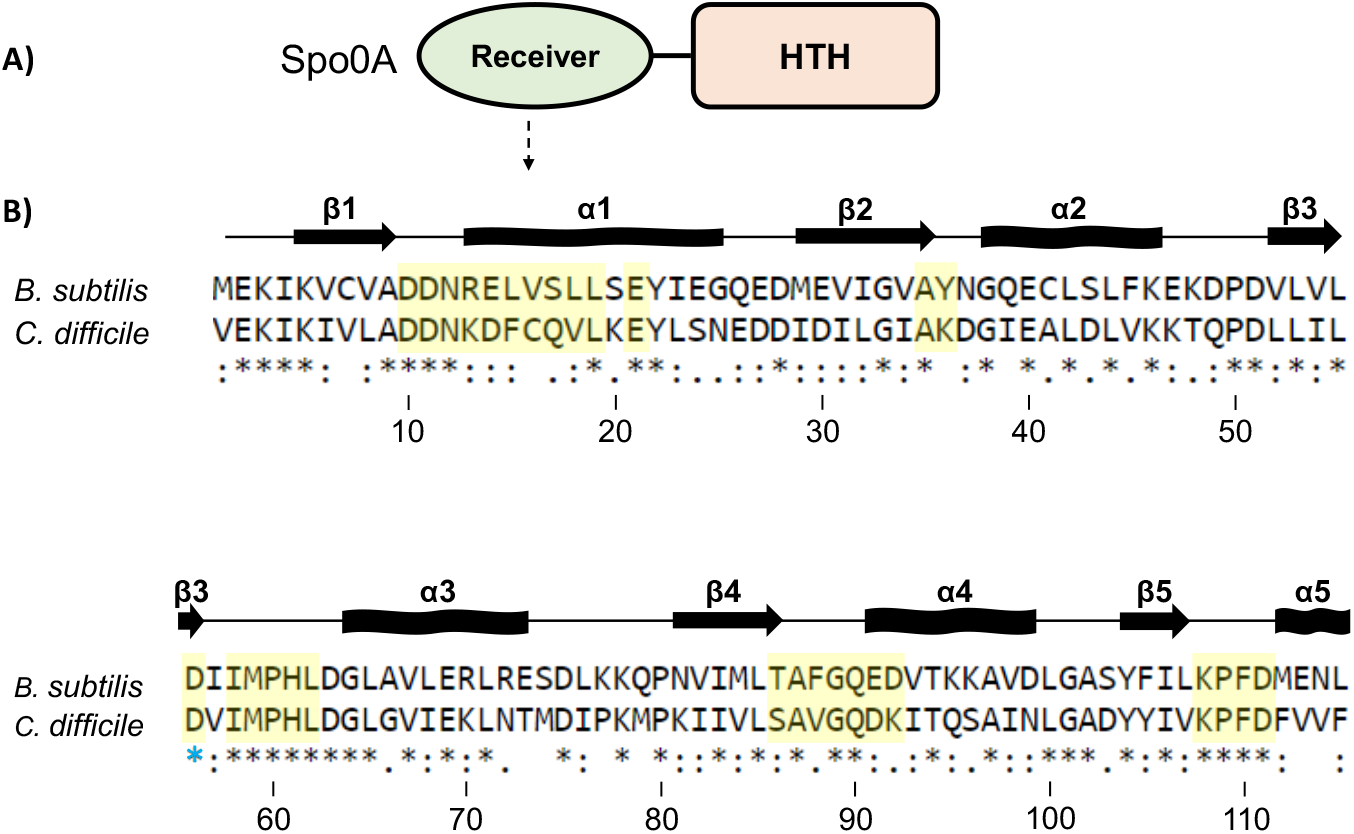
Conservation of the Spo0A receiver domains in *B. subtilis* and *C. difficile*. **A)** Graphic representation of Spo0A domain structure. Functional residues responsible for protein-protein interaction and Spo0A activation are located in the N-terminal receiver domain. The C-terminal region of Spo0A encompasses a helix-turn-helix (HTH) DNA-binding domain. **B)** Alignment of the amino acid sequences of the Spo0A receiver domains for *B. subtilis str. 168* (BSU_24220, top) and *C. difficile* 630 (CD630_12140, bottom). Residues important for *B. subtilis* Spo0F and Spo0A activity that were chosen for mutation in *C. difficile* Spo0A are highlighted in yellow. The blue star (*) is the conserved site of phosphorylation in *B. subtilis*; a corresponding aspartate residue is conserved in *C. difficile* Spo0A. Alignment performed using Clustal Omega. Arrows represent beta sheets, and waved rectangles represent alpha helices.

### Conserved amino acid residues impact Spo0A function in *C. difficile*

To determine the functional significance of the individual mutant Spo0A proteins, the ability for these proteins to complement sporulation when expressed in a *C. difficile spo0A* mutant was assessed. The sporulation frequencies for the mutant *C. difficile spo0A* alleles tested are displayed in **Table 1**. The corresponding *B. subtilis* Spo0A amino acid residue location and the functional significance of each site-directed mutant are also included for reference (**Table 1**).

**Table 1.** Sporulation frequencies of *C. difficile* Spo0A site-directed mutants.

| *C. difficile*<br>Spo0A allele | Spo0A<br>region | Corresponding<br>*B. subtilis*<br>Spo0A residue | Corresponding<br>*B. subtilis*<br>Spo0F residue | Spo0A function in *B. subtilis* | Spo0F function in *B. subtilis* | Average<br>sporulation<br>frequency (%)<sup>a,c</sup> | Mutant<br>phenotype in<br>*B. subtilis*<sup>b</sup> |
| --- | --- | --- | --- | --- | --- | --- | --- |
| Wildtype | - | - | - | Interaction with Spo0B, Spo0E | Interaction with KinA, KinB, KinC, KinD, and KinE, interaction with Spo0B | 12.1±1.0 | - |
| *spo0A::erm* | - | - | - | - | - | 0.00 | - |
| D10A | β1-α1 | D10 | D10 | Forms aspartyl pocket, Spo0A dimerization, divalent cation binding, predicted Spo0E interaction [4, 23, 34, 65] | Forms aspartyl pocket, divalent cation binding [54] | 0.0002±0.0001** | n.d. |
| D11A | β1-α1 | D11 | D11 | Forms aspartyl pocket, Spo0A dimerization, divalent cation binding, predicted Spo0E interaction [4, 23, 34, 65] | Forms aspartyl pocket, divalent cation binding [54] | 0.00055±0.0004* | n.d. |
| N12A | β1-α1 | N12 | Q12 | Interaction with Spo0E, N12K is resistant to hyperactive Spo0E, predicted Spo0B interaction [10, 26, 31-33, 65] | Interaction with Spo0B, inferred interaction with KinA [10, 11, 37, 66] | 14.8±1.86 | **Spo0A gain-of-function** |
| K13A | α1 | R13 | Y13 | n.d. | Interaction with RapA and RapB, Y13S is resistant to RapB [67-70] | 19.1±3 | **Spo0F gain-of-function** |
| D14A | α1 | E14 | G14 | E14A allows direct interaction with KinC, resistant to hyperactive Spo0E [26, 31] | Interaction with Spo0B, and RapB [10, 68-70] | 49.7±6.5* | Spo0A gain-of-function |
| F15A | α1 | L15 | I15 | n.d. | Interaction with KinA, Spo0B, and RapB [10, 11, 68-70] | 23.1±1.7 | **Spo0F decreased sporulation** |
| C16A | α1 | V16 | R16 | n.d. | Interaction with KinA, Spo0B, and RapB [10, 11, 68] | 0.7±0.3* | Spo0F decreased sporulation |
| Q17A | α1 | S17 | I17 | n.d. | Interaction with RapB [11, 66, 68-70] | <LOD** | **Spo0F gain-of-function** |
| V18A | α1 | L18 | L18 | n.d. | Interaction with KinA, Spo0B, RapB [10, 11, 68-70] | <LOD** | Spo0F decreased sporulation |
| L19A | α1 | L19 | L19 | n.d. | Inferred structural importance [11] | 6.4±1.5 | Spo0F decreased sporulation |
| E21A | $\alpha 1$ | E21 | E21 | Inferred to interact with Spo0B [10] | Interaction with Spo0B, inferred interaction with KinA [10, 37] | 0.01±0.01** | n.d. |
| A35S | $\beta 2$ | A35 | A33 | n.d. | n.d. | 0.15±0.1* | n.d. |
| K36A | $\beta 2-\alpha 2$ | Y36 | A34 | | Interaction with Spo0B [10] | 26.4±5.7 | n.d. |
| D56A | $\beta 3$ | D56 | D54 | Site of phosphorylation by Spo0B, forms aspartyl pocket, Spo0A dimerization, predicted to interact with Spo0E [4, 23, 34, 65] | Site of phosphorylation by KinA, KinB, KinC, KinD, and KinE,[25], forms aspartyl pocket [4, 23, 34, 54] | 0.008±0.001** | Spo0A |
| I58A | $\beta 3-\alpha 3$ | I58 | K56 | Stabilizes aspartyl pocket, Spo0A dimerization [23] | Interaction with KinA, Spo0B, and RapB, stabilizes aspartyl pocket [10, 11, 54, 68] | 2±0.9 | Spo0F decreased sporulation |
| M59A | $\beta 3-\alpha 3$ | M59 | I57 | n.d. | Interaction with KinA [11] | 0.006±0.001** | Spo0F decreased sporulation |
| P60A | $\beta 3-\alpha 3$ | P60 | P58 | Interaction with Spo0E, P60S is resistant to hyperactive Spo0E; active without phosphorelay [26, 31] | Interaction with Spo0B [10] | <LOD** | **Spo0A gain-of-function** |
| H61A | $\beta 3-\alpha 3$ | H61 | G59 | n.d. | Interaction with Spo0B [10] | 0.6±0.2* | n.d. |
| L62A | $\beta 3-\alpha 3$ | L62 | M60 | Interaction with Spo0E, L62P is resistant to hyperactive Spo0E [26] | M60A results in reduced *spoII*G transcription [10] | 11.5±1.8 | **Spo0A gain-of-function** |
| S86A | $\beta 4$ | T86 | T82 | Stabilizes phosphorylation of active site [45] | Interaction with KinA, interaction with RapH, interaction with Spo0B [11, 36, 37, 71] | 0.1±0.1* | Spo0F decreased sporulation |
| A87S | $\beta 4$ | A87 | A83 | n.d. | Interaction with Spo0B [10] | 0.01±0.01** | n.d. |
| V88A | $\beta 4-\alpha 4$ | F88 | Y84 | Interaction with Spo0E; F88L is resistant to hyperactive Spo0E [26] | Interaction with Spo0B, KinA, RapH, Y84A is resistant to RapB, RapH [11, 36, 68] | 17.9±1.0 | **Spo0F reduced sporulation** |
| G89A | $\beta 4-\alpha 4$ | G89 | G85 | Inferred to interact with Spo0B [10] | Interaction with Spo0B [10] | 5.6±2.1 | n.d. |
| Q90A | $\beta 4-\alpha 4$ | Q90 | E86 | Q90R allows for direct interaction with KinC, and resistant to hyperactive Spo0E [32, 33] | Interaction with Spo0B, interaction with KinA [10, 11] | 59.3±10.7* | **Spo0F reduced sporulation, Spo0A gain-of-function** |
| D91A | $\alpha 4$ | E91 | L87 | n.d. | Interaction with Spo0B, interaction with KinA [10, 11] | 37.8±12.5 | **Spo0F reduced sporulation** |
| K92A | $\alpha 4$ | D92 | D88 | D92Y is resistant to hyperactive Spo0E and is functional without phosphorelay [25, 31] | n.d. | 33±2.7* | Spo0A gain-of-function |
| K108A | $\beta 5$ | K108 | K104 | Stabilizes aspartyl pocket, Spo0A dimerization, predicted Spo0E and Spo0B interaction [10, 23, 65] | Interaction with Spo0B, stabilizes aspartyl pocket, inferred interaction with KinA [10, 37, 54] | 0.6±0.2* | n.d. |
| P109A | $\beta 5$-$\alpha 5$ | P109 | P105 | Spo0E and Spo0B interaction [10, 65] | Interaction with Spo0B [10] | 76.9±9.3* | n.d. |
| F110A | $\beta 5$-$\alpha 5$ | F110 | F106 | Predicted Spo0E interaction [65] | Interaction with Spo0B [10] | 17.9±2.1 | **Spo0F reduced sporulation** |
| D111A | $\beta 5$-$\alpha 5$ | D111 | D107 | Predicted Spo0E interaction [65] | Interaction with Spo0B, inferred interaction with KinA [10, 37] | 7.6±0.9 | n.d. |

<sup>a</sup> \*, P = > .05; \*\*, P = > .01
<sup>b</sup> *B. subtilis* phenotype that differs from *C. difficile* phenotype noted in bold
<sup>c</sup> Spo0A site-directed mutant sporulation frequency where protein was undetectable by western blot is underlined
n.d., not determined

The strains containing the D14A, Q90A, K92A, and P109A Spo0A site-directed mutants exhibited significantly increased sporulation compared to the control strain expressing wild-type *spo0A* allele (**Figure 3A, 3C**). The strains containing the F15A, K36A, and D91A Spo0A site-directed mutants also displayed increased sporulation but were not statistically significant (**Table 1**). The *C. difficile* Spo0A Q90A gain-of-function sporulation phenotype was similar to the increased sporulation phenotype observed with the *B. subtilis* Spo0A Q90R mutant, which facilitates interaction with the activating protein KinC [31-33]. The increased sporulation phenotype displayed by the *C. difficile* Spo0A D14A mutant was similar to the sporulation phenotype observed when the orthologous *B. subtilis* Spo0A residue E14 is mutated [26, 31]. The *B. subtilis* Spo0A E14A mutant confers resistance to hyperactive Spo0E, resulting in increased Spo0A phosphorylation and activity [26, 31]. The gain of function phenotype of the *C. difficile* D14A mutant suggests that this residue may also be important for recognition by Spo0E in *C. difficile* [26, 31]. The *B. subtilis* Spo0F residues G14, L87, and P105 are all important for positively influencing sporulation through interaction with Spo0B (**Table 1**), yet the corresponding *C. difficile* site-directed mutants (Spo0A D14A, D91A, and P109A) all exhibited increased sporulation, suggesting that these residues serve a divergent role in *C. difficile* Spo0A activation [10].

**Figure 3.**
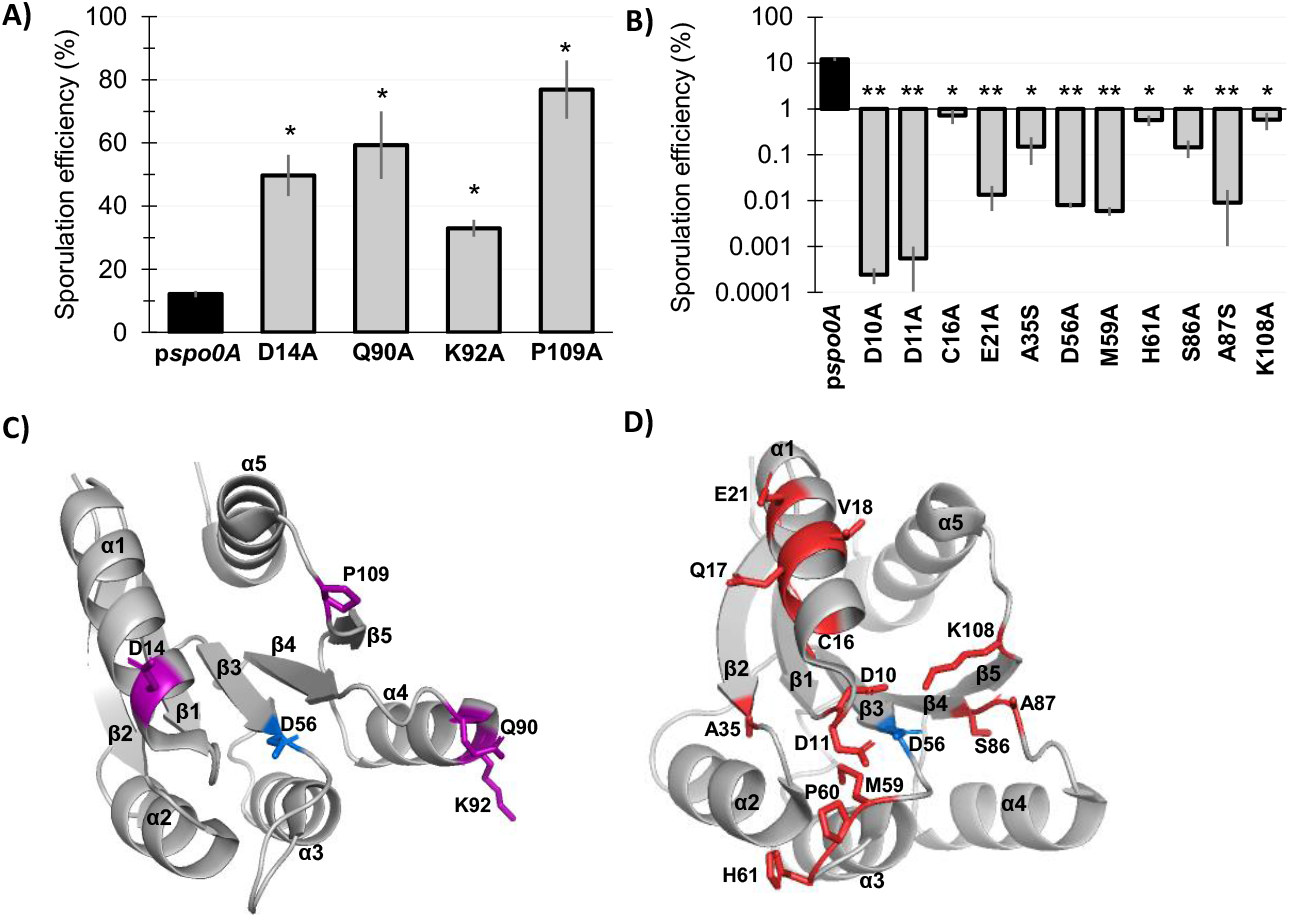
Mutagenesis of conserved Spo0A residues results in both increased and decreased *C. difficile* sporulation frequency. **A)** Ethanol-resistant spore formation of 630Δ*erm spo0A* p*spo0A* (MC848) expressed on a plasmid compared to the Spo0A site-directed mutants D14A (MC1671), Q90A (MC1712), K92A (MC1185), and P109A (MC1621) with increased sporulation frequency. **B)** Ethanol-resistant spore formation of 630Δ*erm spo0A* p*spo0A* (MC848) expressed on a plasmid compared to the Spo0A site-directed mutants D10A (MC1618), D11A (MC1703), C16A (MC1057), E21A (MC1058), A35S (MC1059), D56A (MC849), M59A (MC1184), H61A (MC1036), S86A (MC1846), A87S (MC1061), and K108A (MC1064) with decreased sporulation frequency, displayed on log_10_ scale. Sporulation assays were performed independently at least four times. Statistical significance was determined using Kruskal-Wallis test and uncorrected Dunn’s test (*, P = > 0.05; **, P = > 0.01). **C)** 3D structure of Spo0A with residues (highlighted purple) that cause increased sporulation when mutated, orientated around the activation site (D56, highlighted blue). **D)** 3D structure of Spo0A with residues (highlighted red) that reduce sporulation when mutated, orientated around the active site (D56, highlighted blue). Spo0A PDB code 5WQ0, edited in PyMOL (The PyMOL Molecular Graphics System, Version 2.4.0 Schrödinger, LLC).

Conversely, 15 of the 30 Spo0A site-directed mutants had reduced sporulation, representing a much larger proportion of the mutants assessed (**Table 1, Figure 3B, 3D**). Expression of eleven of the mutant *spo0A* alleles resulted in significantly reduced sporulation: D10A, D11A, C16A, E21A, A35S, D56A, M59A, H61A, S86A, A92S, and K108A. Spo0A Q17A, V18A, and P60A demonstrated sporulation frequencies below the limit of detection (>0.0002%); however, through western blotting we found these Spo0A site-directed mutants were not stably produced (**Table 1, Supplemental Figure 2**).

The *C. difficile* Spo0A mutants that demonstrate a loss-of-function sporulation phenotype may represent amino acid residues that facilitate direct interactions with positive regulators of sporulation. The *C. difficile* Spo0A C16A and E21A mutations are located within the α1 region (**Figure 3D**). The *B. subtilis* Spo0F equivalents, R16 and E21, promote direct interactions with KinA and Spo0B [11], while *B. subtilis* Spo0A E21 is also expected to interact with Spo0B [10]. Altogether, these data suggest that *C. difficile* C16 and E21 coordinate the interaction with a positive regulator of Spo0A activity. The *B. subtilis* Spo0A residues D10, D11, I58, and K108 form the aspartyl pocket and are important for Spo0A homodimerization, which is necessary for DNA-binding activity [23, 34]. Mutation of the *C. difficile* Spo0A equivalent residues D10, D11, and K108 all produced severe sporulation defects. The *C. difficile* Spo0A I58A mutant had decreased sporulation, although these results were not statistically significant.

The aspartate residue at position 56 of *B. subtilis* Spo0A serves as the phosphorylation site and is critical for sporulation, consistent with findings for the conserved aspartate residue in other species’ Spo0A orthologs [4, 14-17, 35]. As expected, mutation of the predicted *C. difficile* Spo0A phosphorylation site (D56A) resulted in dramatically reduced sporulation (>1000-fold decrease, **Table 1**), suggesting that this aspartate residue is required for *C. difficile* Spo0A phosphorylation and activation. *C. difficile* Spo0A I58, M59, and H61 are located immediately adjacent to the phosphorylation site in the open face between β3-α3 (**Figure 3D**). Mutation of the *B. subtilis* Spo0F K56 residue results in a loss-of-function phenotype, and several residues in this region facilitate *B. subtilis* Spo0A and Spo0F interactions with kinases or phosphatases [10, 11]. These data correspond with the low sporulation frequencies of the orthologous *C. difficile* Spo0A I58A, M59A, and H61A mutants (**Figure 3B**), suggesting that this region functions similarly in *C. difficile*. The β4 region of *B. subtilis* Spo0A is important for phosphotransfer between Spo0F or Spo0B. The *C. difficile* Spo0A S86 and A87 residues are located at the C-terminal end of β4 (**Figure 3D**), and site-directed mutagenesis of these residues significantly reduced sporulation frequency (**Table 1**), suggesting that the β4 region is likewise important for phosphotransfer to *C. difficile* Spo0A [10, 36]. Additionally, the *B. subtilis* Spo0F residue T82 is equivalent to *C. difficile* Spo0A S86, and is involved in stabilizing the phosphorylation of Spo0F [37]. Since both threonine and serine have polar side chains, *C. difficile* S86 may also facilitate phosphorylation of the Spo0A active site. Finally, the *C. difficile* Spo0A A35 residue is conserved in both *B. subtilis* Spo0A (A35) and Spo0F (A33), though the function of these residues in Bacilli have not been determined.

The Spo0A mutants N12A, K13A, L19A, L62A, F110A, and D111A had sporulation frequencies that were comparable to the wildtype Spo0A allele. N12, K13 and L19 appear to be dispensable for sporulation, even though residues located in this region are important for interaction of *B. subtilis* Spo0A and Spo0F with both positive and negative regulators (**Table 1**) [11, 31]. However, the Spo0A L19A mutant exhibited a translucent plate morphology on sporulation agar, suggesting some functional importance in other physiological processes (**Table 2**). This result is not surprising given the pleiotropic effects Spo0A displays in *C. difficile* and other species [8, 17, 38-44]. Mutation of the *Bacillus* Spo0A and Spo0F residues that are comparable to *C. difficile* Spo0A L62 result in gain-of-function phenotypes, but was not important for *C. difficile* Spo0A activity (**Table 1**). The *C. difficile* Spo0A residues F110A and D111A are located at the open face of β5-α5 in a motif (KPFD) that is highly conserved in the CheY superfamily of response regulators [10, 45]. Our data indicate that this region is also important for *C. difficile* Spo0A regulation, as the K108A mutant had decreased sporulation and the P109A mutant displayed increased sporulation (**Table 1**). While the F110A or D111A mutants did not affect sporulation, mutation of these residues produced a translucent and crushed plate morphologies, respectively (**Table 2**), suggesting that they impact Spo0A function.

**Table 2.** Spo0A site-directed mutant morphology and growth phenotypes.

| **Spo0A mutant** | **Morphology phenotype** |
| --- | --- |
| D14A | Mucoidal |
| F15A | Mucoidal |
| C16A | Mucoidal |
| Q17A | Poor growth |
| L19A | Translucent |
| E21A | Mucoidal, poor growth |
| A35S | Mucoidal, poor growth |
| H61A | Mucoidal, poor growth |
| A87S | Mucoidal, poor growth |
| Q90A | Mucoidal |
| D91A | Mucoidal |
| P109A | Mucoidal |
| F110A | Translucent |
| D111A | Crushed |

### Altered growth and morphology of Spo0A mutants

Fourteen of the mutated *spo0A* alleles produced phenotypes that impacted growth in BHIS broth and growth and morphology on sporulation agar (**Table 2**). The most commonly observed phenotype was a stringy, mucoidal morphology that was observed for ten of the mutants after 24 h growth on sporulation plates (Spo0A D14A, F15A, C16A, E21A, A35S, H61A, A87S, Q90A, D91A, and P109A). The mucoidal phenotype was observed in both hyper- and hyposporulating strains, indicating that mucoidy is not directly correlated with the sporulation outcome. The Spo0A L19A and F110A mutants produced flat, translucent lawns, but sporulation was not affected in either mutant background. Similarly, the Spo0A D111A mutant did not affect sporulation, but produced a rigid, crushed lawn morphology. Strains expressing *spo0A* Q17A, E21A, A35S, H61A, and A87S exhibited poor growth in BHIS liquid compared to expression of the wildtype *spo0A* (data not shown). The Spo0A mutants with poor growth all had reduced sporulation (**Table 1**). However, only 5 of the 14 hyposporulating mutants grew slowly, indicating that defects in Spo0A that reduce sporulation do not necessarily retard growth.

### *C. difficile* Spo0A requires phosphorylation of the conserved aspartate for activation

In *B. subtilis*, Spo0A is phosphorylated at the conserved aspartate residue D56, which is required for activation [34, 46]. In the activated state, Spo0A homodimerizes and binds to specific DNA sequences, or “0A boxes”, to regulate Spo0A-dependent gene expression [47, 48]. Sequence comparison to *B. subtilis* Spo0A and other response regulators implicated D56 as the conserved site of *C. difficile* Spo0A phosphorylation and activation. The *C. difficile* Spo0A D56A site-directed mutation also dramatically reduced sporulation, further supporting the necessity of this residue for activity (**Figure 3B**). To determine if *C. difficile* Spo0A is also phosphorylated at the conserved aspartate residue, we isolated total protein from strains expressing either p*spo0A*-3XFLAG 3x-FLAG-Spo0A or p*spo0A*-D56A-3XFLAG and separated phosphorylated and unphosphorylated Spo0A species using phos-tag SDS-polyacrylamide gel electrophoresis followed by western blotting with an α-FLAG antibody [49-51] (**Figure 4**). In the phos-tag assay, higher molecular weight bands that are present in the unheated sample but absent in the heated sample represent phosphorylated protein, as phosphoryl groups are heat-labile. In the strain expressing wild type *spo0A*, two bands were observed in the unheated sample, with the upper band denoting phosphorylated Spo0A and the lower band corresponding to unphosphorylated Spo0A. In contrast, the Spo0A D56A mutant displayed only the lower, unphosphorylated band in both the unheated and heated samples, indicating that D56 is the primary site of phosphorylation. Together, the sporulation defect of the D56A mutant and the absence of Spo0A phosphorylation demonstrate that D56 is the primary site of Spo0A phosphorylation.

**Figure 4.**
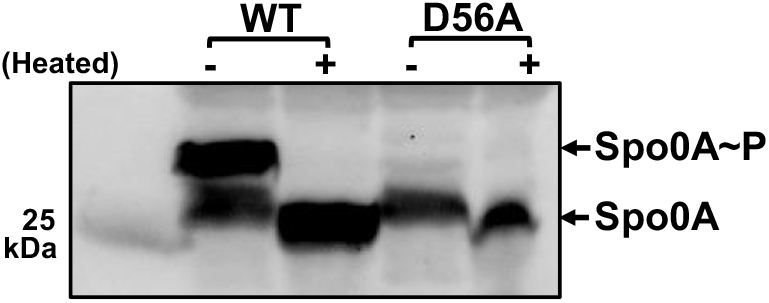
The conserved aspartate residue of *C. difficile* Spo0A is phosphorylated. Anti-FLAG western blot after phos-tag gel separation of unphosphorylated and phosphorylated Spo0A (Spo0A∼P) species in 630Δ*erm spo0A* p*spo0A-*3XFLAG (MC1003) and 630Δ*erm spo0A* p*spo0A* D56A*-*3XFLAG (MC1690) grown on sporulation agar for 12 h. The molecular weight marker (25 kDa) is indicated on the left of the panel. Shown is a representative blot from one of four independent experiments.

### Residues necessary for Spo0A dimerization in other species have conserved functions in *C. difficile*

Residues that are important for Spo0A homodimerization were previously identified in Bacilli [23, 34, 45]. The residues of *C. difficile* Spo0A that facilitate dimerization have not been characterized; however, *C. difficile* Spo0A contains five residues that are identical to those involved in dimerization in *B. subtilis* and other aerobic spore-formers: D10, D11, D56, I58, and K108. The alanine mutants of these five residues all produced defects in sporulation, indicating that they are important for Spo0A function. To test if these residues are involved in Spo0A dimerization *in vivo*, we performed split-luciferase reporter assays. Here, luciferase enzyme is fragmented into either a SmBit or LgBit subunit and fused to a gene(s) of interest to test for protein-protein interaction [52, 53]. We constructed C-terminal fusions of the SmBit and LgBit luciferase subunits to the wildtype, D10A, D11A, D56A, I58A, and K108A mutant *spo0A* alleles. All five site-directed mutants had less activity than the wildtype Spo0A fusions, with the D10A and D56A alleles exhibiting significantly less output (**Figure 5A, Supplemental Table 1**), indicating that these Spo0A site-directed mutations reduce the ability for these mutant proteins to form homodimers. The D10, D11, I58, and K108 residues are all oriented around the D56 activation site (**Figure 5B**), further supporting the importance of these residues for Spo0A homodimerization [23]. These results demonstrate that the functional residues that are involved in Bacilli Spo0A dimerization are also important for *C. difficile* Spo0A dimerization.

**Figure 5.**
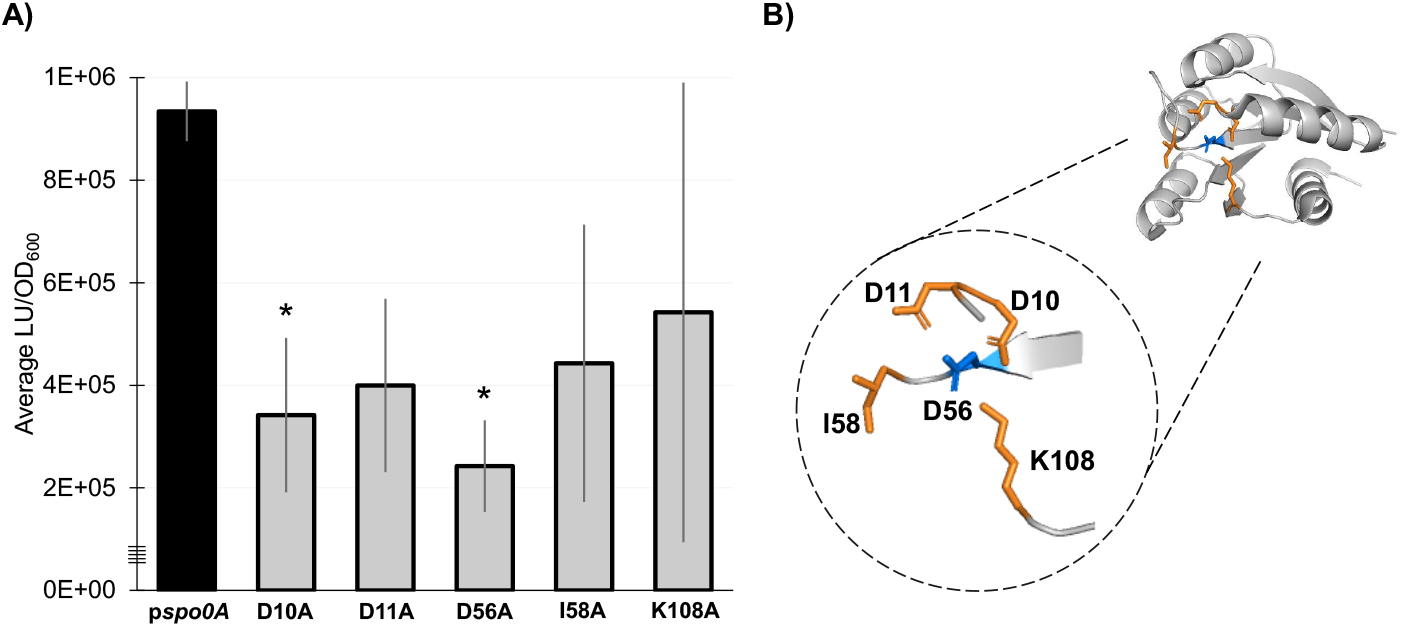
Residues necessary for Spo0A dimerization in other Firmicutes are functionally conserved in *C. difficile*. **A)** Split-luciferase activity in strains 630Δ*erm spo0A* p*spo0A* (MC1906) and the Spo0A site-directed mutants D10A (MC2001), D11A (MC2002), D56A (MC2003), I58A (MC2004), and K108A (MC2005) fused to SmBit and LgBit fragments after cultures were grown in 70:30 sporulation broth to OD_600_ = 0.8 – 0.9 and induced with anhydrous tetracycline (ATc) for 1 h. Average luminescence outputs are normalized to optical densities (LU/OD_600_). Error bars represent the standard deviation of three independent experiments (*, P = < 0.05) as determined by a one-way ANOVA with Dunnett’s multiple comparisons test. **B)** 3D structure of Spo0A with the residues that form the aspartyl pocket and facilitate dimerization highlighted orange near the site of activation (D56, highlighted blue). Spo0A PDB code 5WQ0, edited in PyMOL (The PyMOL Molecular Graphics System, Version 2.4.0 Schrödinger, LLC).

## DISCUSSION

In this study, we employed alanine-scanning mutagenesis to define the regions of the *C. difficile* Spo0A receiver domain that are important for regulation of sporulation. Altogether, we examined the ability of *C. difficile* Spo0A to initiate sporulation through mutational analysis of 30 residues located within 10 different regions of the Spo0A receiver domain secondary structure (**Figure 2**). The results demonstrated that mutation of many residues that influence *B. subtilis* Spo0A and Spo0F activation also have profound effects on *C. difficile* Spo0A function, even though few of the interacting partner proteins are conserved between these species. We also established Spo0A residues that are important for homodimerization and found altered growth and morphology phenotypes by mutating the receiver domain of Spo0A.

The receiver domain of the *C. difficile* Spo0A shares 47% identity with the *B. subtilis* Spo0A and 30% identify with *B. subtilis* Spo0F, but the protein architectures of the receiver motifs are highly conserved. By probing the function of conserved regions and residues that have been implicated in protein interaction in Bacilli, we demonstrate conservation of sporulation phenotypes in many *C. difficile* Spo0A residues relative to *Bacillus* Spo0F and Spo0A (**Table 1**) [10, 11, 23, 25, 28, 32]. As in Bacilli, we found that the receiver domain α-helices and the β1-α1, β3-α3, β4-α4, and β5-α5 open faces are all important for *C. difficile* Spo0A activity (**Table 1**) [10, 11, 23]. The majority of residues that were mutated in this study that produced major changes in sporulation are orientated on the same face as the site of activation, an effect observed for other response regulator receiver domains (**Figure 3**) [11, 45, 54].

*C. difficile* Spo0A and *B. subtilis* Spo0A perform the same sporulation function, but there are gaps in knowledge about the contribution of specific residues to Spo0A activity in both species. Our data demonstrate that most residues within the receiver domains of *B. subtilis* and *C. difficile* Spo0A proteins have similar impacts on sporulation. However, many of the characterized *B. subtilis* Spo0A site-directed mutants are gain-of-function suppressor mutations that are not alanine substitutions or were characterized in strain backgrounds lacking elements of the phosphorelay [25, 31]. Additionally, many of the described residues that are important for sporulation in *B. subtilis* Spo0F have not been characterized in Spo0A. Our sporulation results in *C. difficile* suggest open questions remain about the function of the following *B. subtilis* Spo0A residues: L15, V16, S17, L18, E21, A35, I58, M59, P60, H61, T86, A87, Q90, E91, D92, K108, and P109. These residues may also be important for Spo0A function in *B. subtilis* and other spore-forming Firmicutes.

The receiver domain of *B. subtilis* Spo0F has been more extensively characterized than Spo0A and more is understood about the impact of specific Spo0F residues on the regulation of sporulation [10, 11, 37]. We found several differences in the sporulation outcomes for mutations in conserved residues of *B. subtilis* Spo0F and *C. difficile* Spo0A, which is not surprising, considering the differences in these species’ sporulation pathways. Mutation of *C. difficile* Spo0A residues K13, F15, Q17, L62, V88, Q90, D91, and F110 resulted in different impacts on sporulation relative to similar mutations in Spo0F (**Table 1**). Some of these residues are important for Spo0F interaction with factors that are not present in *C. difficile*, such as the Rap phosphatases, Spo0B, and the specific sporulation kinases of *Bacillus* (**Figure 1, Table 1**). In particular, the *C. difficile* Spo0A mutants F15A, Q90A, and D91A displayed higher sporulation, while corresponding residues in *B. subtilis* Spo0F (I15A, E86A, L87A) resulted in sporulation defects [11]. *C. difficile* Spo0A V88A and F110A maintained wild type sporulation levels, while *B. subtilis* Spo0F Y84A and F106A resulted in sporulation defects [11]. These results suggest that the importance of these residues is maintained for both proteins, although the interacting partners and the resulting effects on sporulation differ.

Distinct effects on growth and colony morphology were observed for the 14 Spo0A mutants listed in **Table 2**. The growth and morphology phenotypes are likely due to altered Spo0A regulation or function, as we found that deletion of *spo0A* in *C. difficile* does not change growth or morphology under the conditions tested, as previously observed [20, 38, 39]. The mucoidal phenotype observed on sporulation agar was the most commonly observed effect and was found in 10 of the 30 characterized Spo0A mutants. Mucoidy was only observed when the mutants grew for at least 12 hours as a lawn on sporulation agar, suggesting this phenotype is linked to either a facet of sporulation or conditions that facilitate sporulation (data not shown). However, the mucoidal phenotype was present in both hyposporulating and hypersporulating mutants and did not have an obvious impact on the capacity to sporulate. Additional changes in morphology, but not sporulation, were observed in Spo0A L19A, F110A and D111A. The L19A and F110A mutants produced flat, translucent lawns on sporulation agar, and the D111A mutant had a crushed morphology on sporulation agar. Further, the mutants Q17A, E21A, A35S, H61A, and A87S all had poor growth in BHIS broth relative to both the wildtype and *spo0A* mutant, and all had defects in sporulation. To our knowledge, this is the first report that specific Spo0A residues impact colony morphology or growth. While it is unclear why Spo0A mutant alleles would affect morphology or growth, the simplest explanation is that the altered Spo0A alleles can interact with additional partner proteins that control these cellular processes. It remains unknown if the morphology and growth phenotypes of specific Spo0A site-directed mutants are unique to *C. difficile* or if these phenotypes are conserved for Spo0A of other spore-forming Firmicutes.

We found that Spo0A is phosphorylated at the conserved site of activation (D56), and that a D56A mutation results in loss of phosphorylation (**Figure 4**). Interestingly, the D56A mutant does exhibit reduced, but not total loss of sporulation. This could be a result of low Spo0A DNA-binding activity present in unphosphorylated Spo0A. Although all studied Spo0A are regulated by phosphorylation and dephosphorylation, the proteins that directly interact with Spo0A vary considerably within the Clostridia and the Spo0A proteins in these species have diverged (**Supplemental Figure 3A, 3B**). In other Clostridia in which Spo0A regulation has been studied, Spo0A is acted upon directly by orphan histidine kinases or phosphatases to regulate Spo0A activity, and all encode at least one kinase that induces sporulation [14-18, 55, 56]. While a Spo0A-activating kinase has not yet been identified in *C. difficile*, our data confirm that phosphorylation of Spo0A at the conserved site of activation is critical for Spo0A activity.

To our knowledge, this represents the first report on residues important for Spo0A dimerization in *C. difficile* (**Figure 5**) [23, 34]. The fact that the mechanism of dimerization is maintained in *C. difficile* is likely due to the conserved architecture of response regulator receiver domains, defined by (β/α)_5_ folding and functional residues that are orientated near the site of activation [23, 34, 45, 54].

The Bacilli and Clostridia diverged roughly 2.4 billion years ago during the Great Oxidation Event [57]. While the mechanism(s) of *C. difficile* Spo0A regulation remains unclear, we have identified conserved regions of Spo0A that are important for activity. Because the phosphorelay interactions are not retained in *C. difficile*, our results suggest that *C. difficile* Spo0A uses functionally conserved regions for interaction with both positive and negative regulators that are not part of the Bacilli mechanism for Spo0A regulation. Elucidation of the factors that regulate Spo0A in *C. difficile* will provide greater insight on the biology and lifestyle of this clinically important pathogen.

## MATERIALS AND METHODS

### Bacterial strains and growth conditions

Bacterial strains and plasmids used in this study are listed in (**Table 3**). *C. difficile* strains were routinely grown in BHIS broth or on BHIS agar supplemented with 2-5 μg ml^-1^ thiamphenicol (Sigma-Aldrich) as needed [58]. *C. difficile* cultures were supplemented with 0.2% fructose and 0.1% taurocholate (Sigma-Aldrich) to prevent sporulation and induce germination as indicated [30, 58]. *C. difficile* was grown on 70:30 agar to assess sporulation frequency as previously described [30]. *C. difficile* strains were grown in a 37°C anaerobic chamber (Coy) with an atmosphere consisting of 10% H_2_, 5% CO_2,_ and 85% N_2_, as previously described [59]. Strains of *Escherichia coli* were cultured in LB at 37°C [60] and supplemented with 100 μg ml^-1^ ampicillin or 20 μg ml^-1^ chloramphenicol as needed. Kanamycin 100 μg ml^-1^ was used for counterselection of *E. coli* HB101 pRK24 after conjugation with *C. difficile* [61].

**Table 3.**
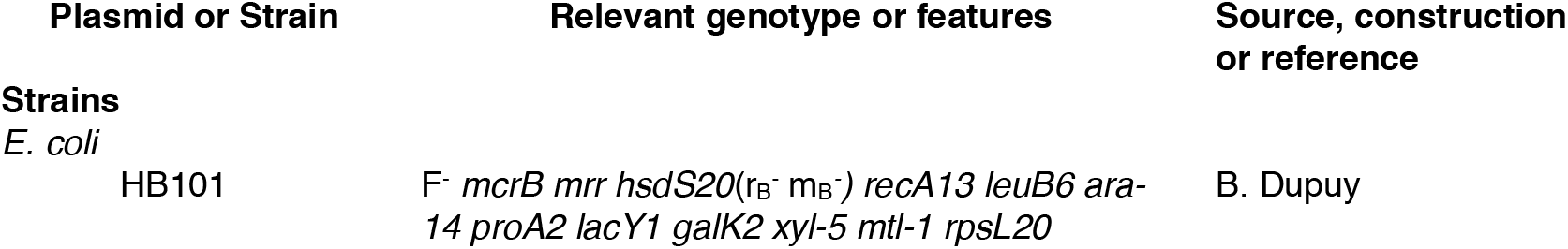
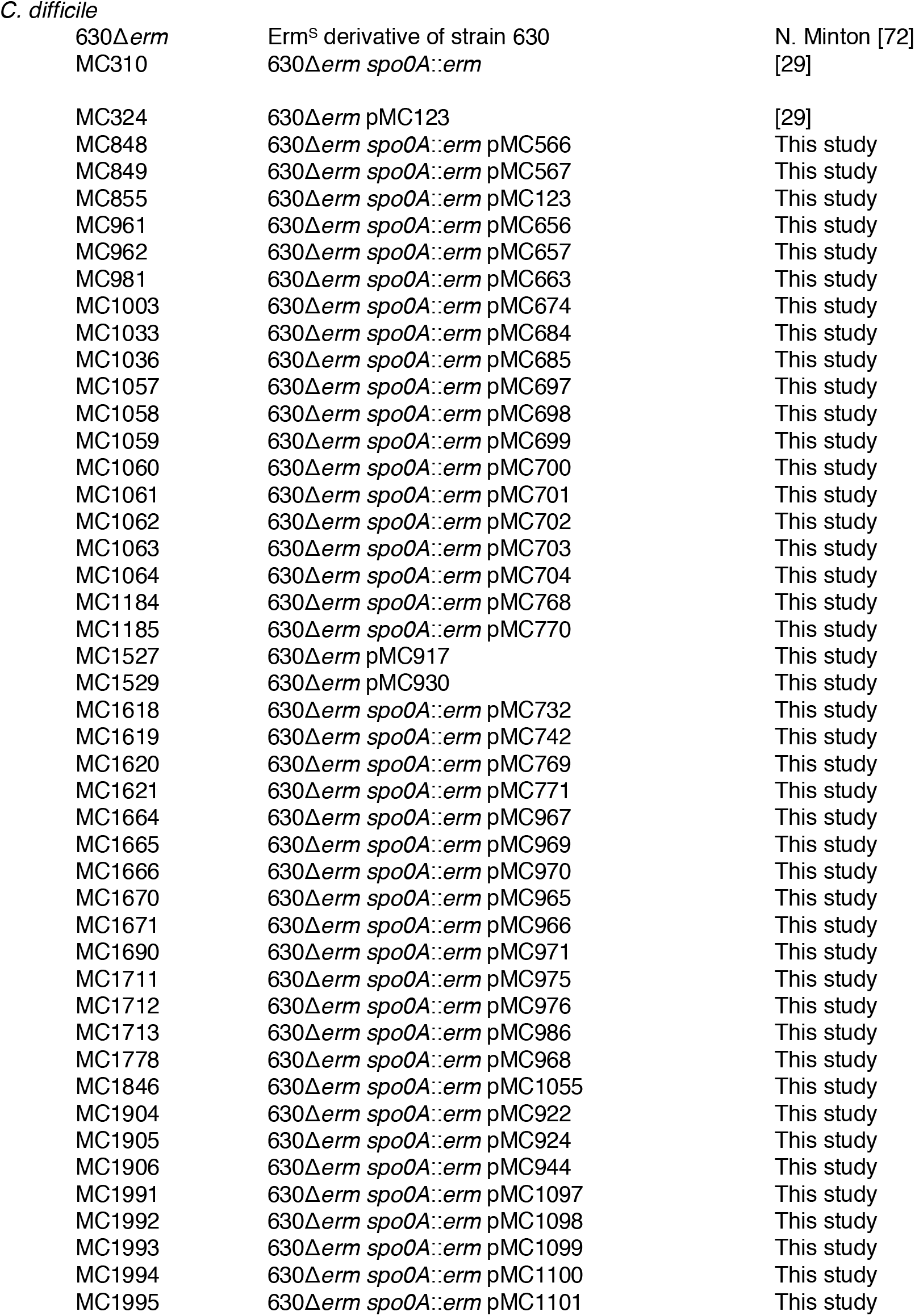
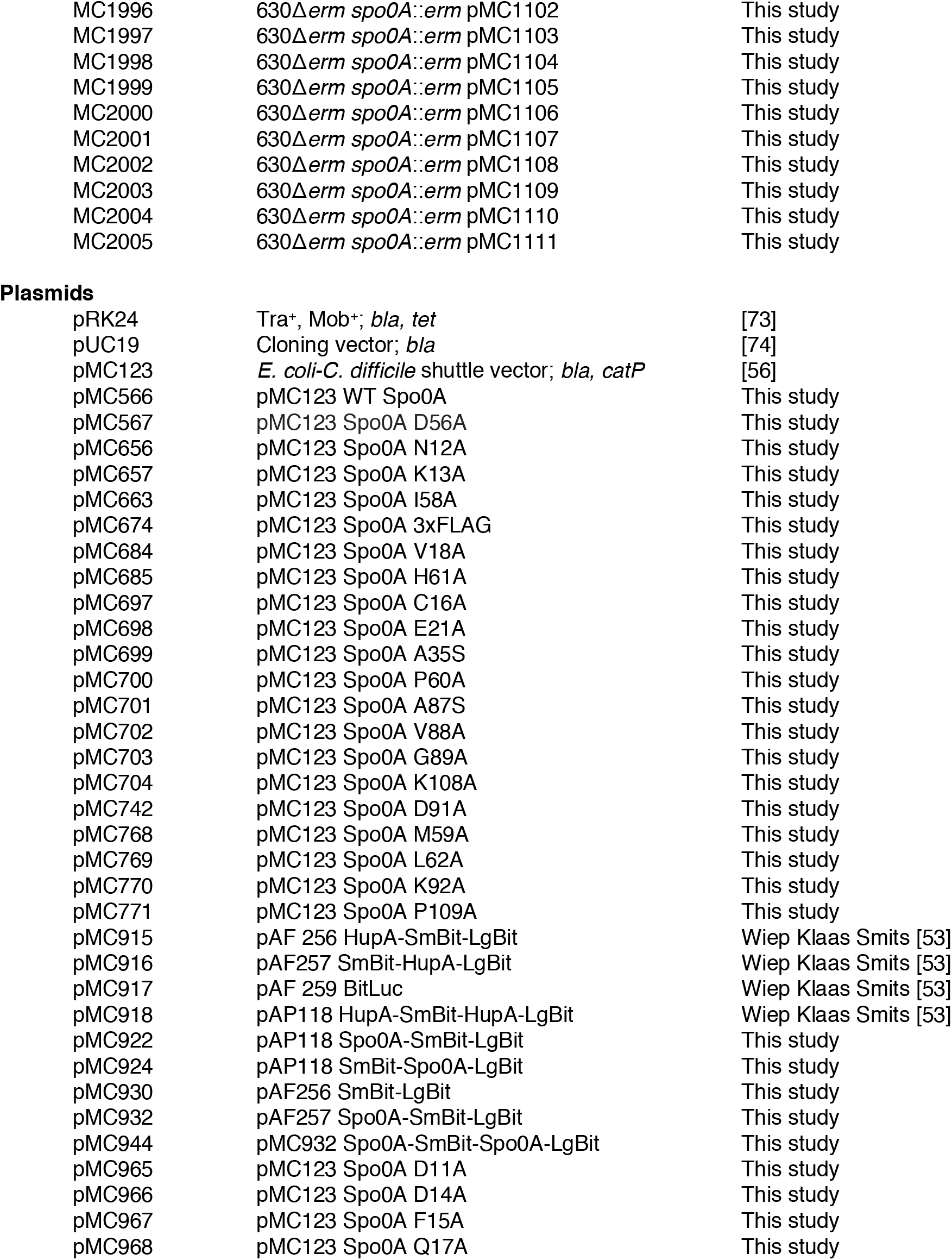
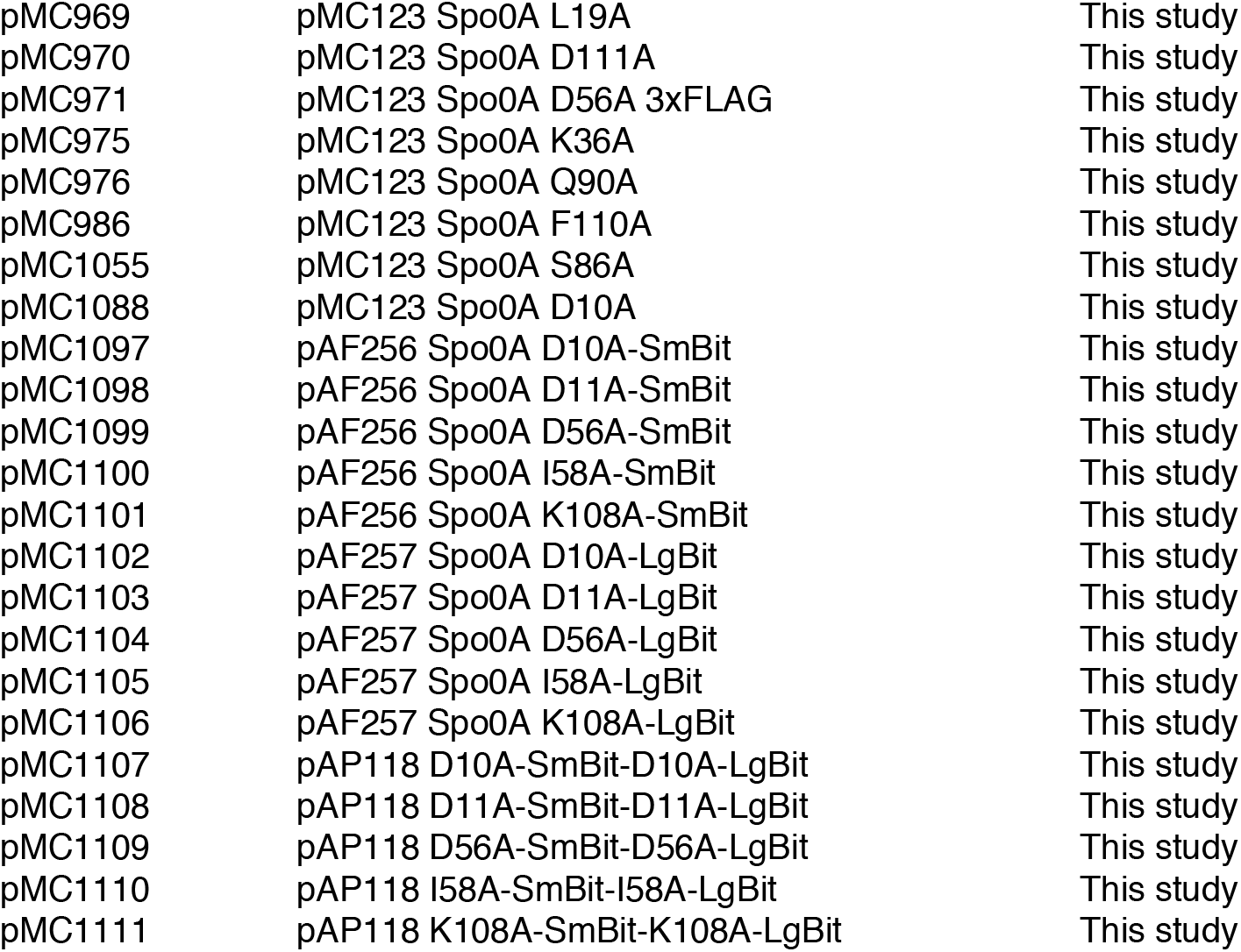
Bacterial Strains and plasmids.

### Strain and plasmid construction

**Table 4** contains oligonucleotides used in this study. *C. difficile* 630 strain (GenBank accession number **<u>AJP10906.1</u>**) was used as a template for primer design and *C. difficile* 630Δ*erm* genomic DNA was used for PCR amplification. *C. difficile* 630Δ*erm* has a known 18 nucleotide duplication outside of the Spo0A receiver domain and was used for strain creation [8]. Strain construction is described in **Supplemental Table 2**.

**Table 4.**
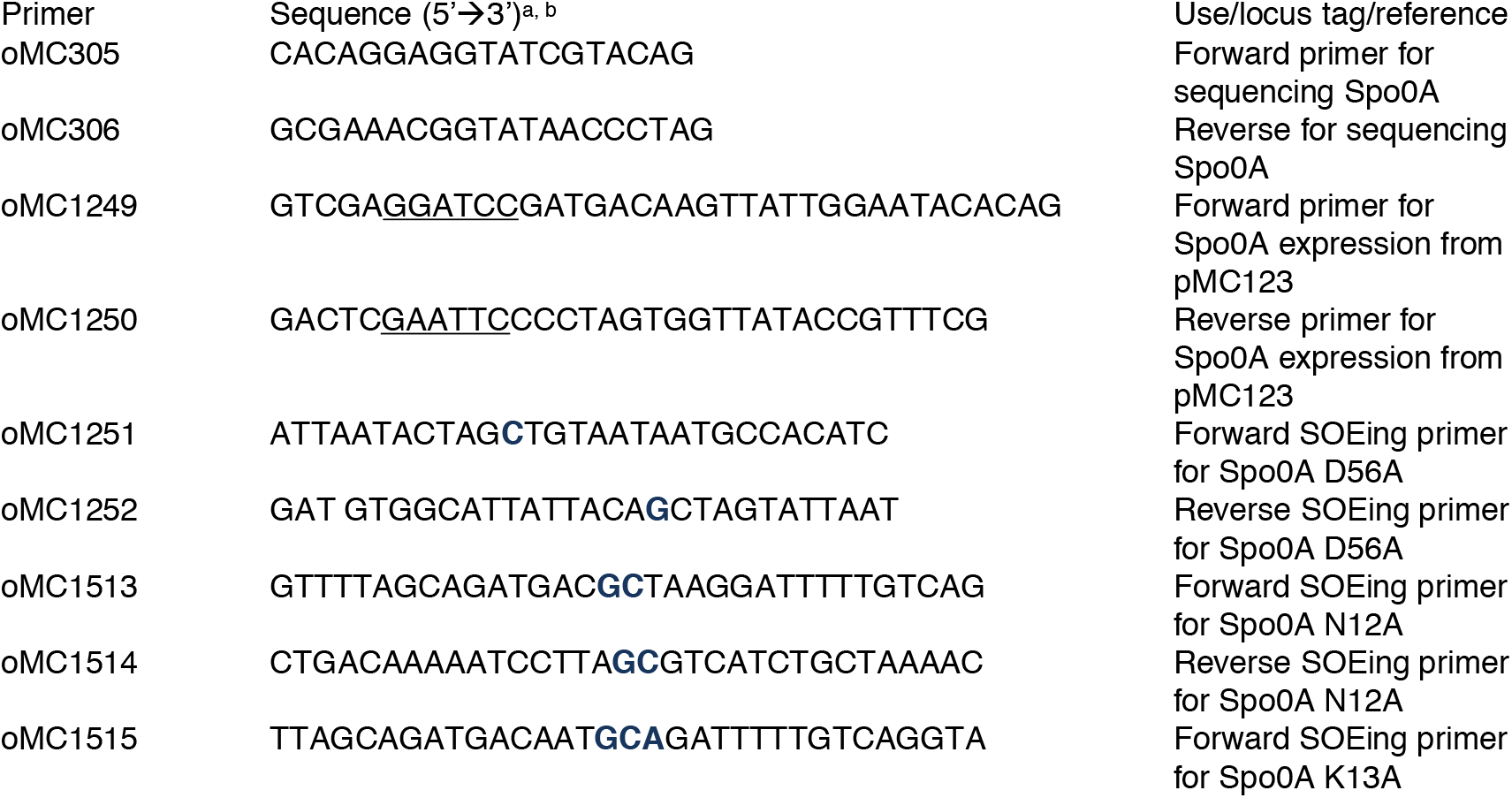
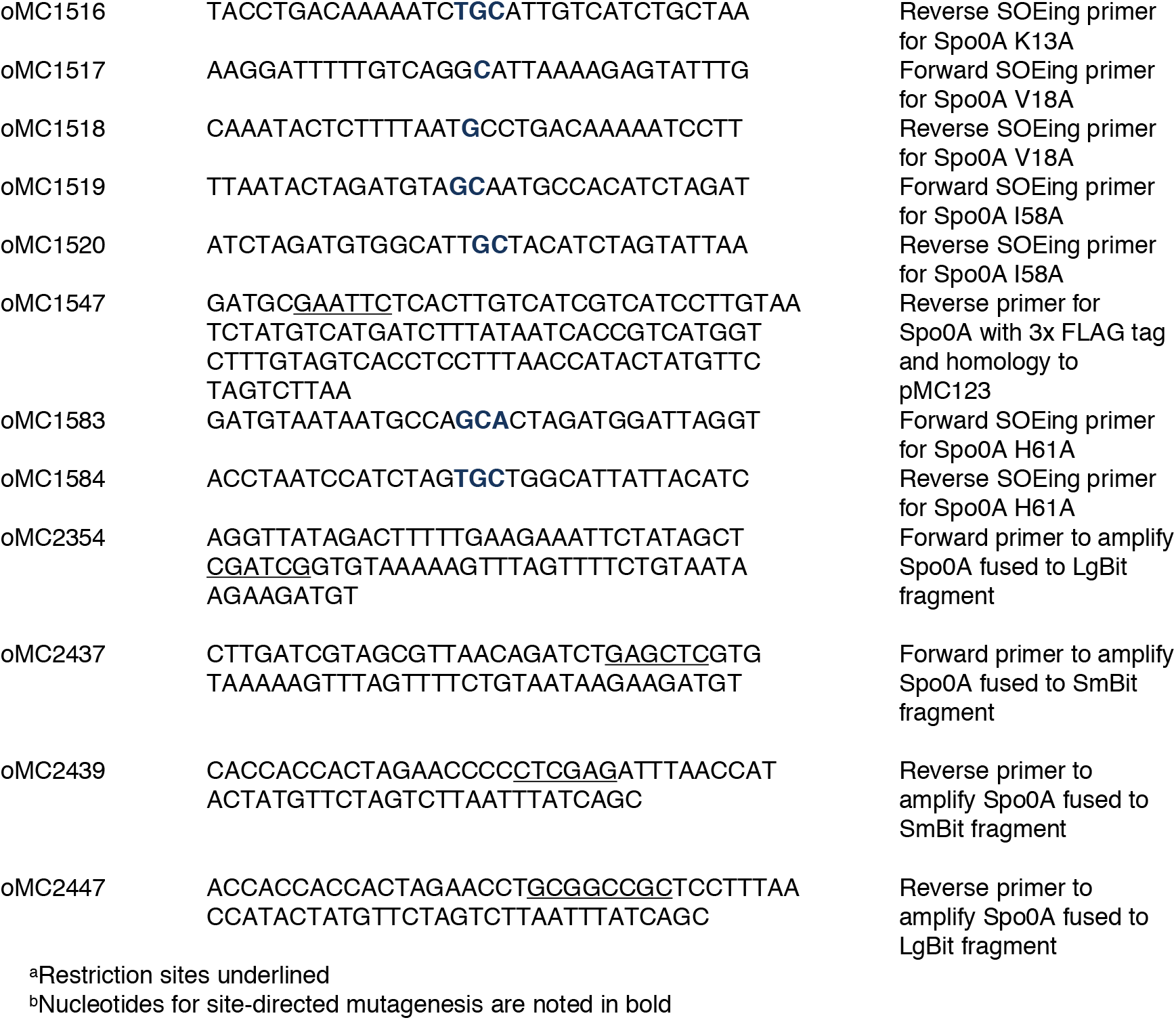
Oligonucleotides.

### Dendrogram

The Spo0A dendrogram rooted *to B. subtilis* Spo0A was created using the MUSCLE Multiple Alignment plugin and Geneious Tree Builder in Geneious Prime v2020.2.2. Spo0A amino acid sequences from *C. difficile* 630 (GenBank accession **<u>AJP10906.1</u>**), *C. perfringens* SM101 (GenBank **<u>CP000312.1</u>**), *C. acetobutylicum* ATCC 824 (GenBank **<u>NC_003030.1</u>**), *A. thermocellus* DSM 1313 (GenBank **<u>NC_017304.1</u>**), *C. botulinum* A str. ATCC 3502 (GenBank **<u>NC_009495.1</u>**), and *B. subtilis* str. 168 (GenBank **<u>NC_000964.3</u>**) were retrieved, aligned, and assembled into a dendrogram. The percentage of identity for each Spo0A protein relative to *B. subtilis* and the heatmap comparing Spo0A percent identities to each species was generated using Geneious Tree Builder in Geneious Prime v2020.2.2. (https://www.geneious.com).

### Sporulation assays

*C. difficile* ethanol resistance sporulation assays were performed on 70:30 sporulation agar supplemented with 2 μg ml^-1^ thiamphenicol for plasmid maintenance, as previously described [62-64]. Following growth on sporulation agar for 24 h, cells were resuspended in BHIS broth to an OD_600_ of 1.0. To determine total vegetative cell counts ml^-1^, cultures were serially diluted in BHIS and plated on BHIS agar with 2 μg ml^-1^thiamphenicol. Concurrently, 0.5 ml of resuspended cells were treated with a mixture of 0.3 ml 95% ethanol and 0.2 ml dH_2_O for 15 min to kill all vegetative cells, then serially diluted in a mixture of 1X PBS and 0.1% taurocholate and plated onto BHIS agar with 2 μg ml^-1^ thiamphenicol and 0.1% taurocholate to enumerate the total number of spores per ml. After 48 h growth, CFU were calculated and the sporulation frequency was determined as the number of spores that germinated following ethanol treatment divided by the total number of spores and vegetative cells [62]. A *spo0A* mutant complemented with wild-type *spo0A* driven from its native promoter on a plasmid was used as a positive control (MC848), and a *spo0A* null mutant containing the empty vector was used as the negative control (MC855). Statistical analyses were performed using the Welch’s ANOVA with Dunnett’s multiple comparisons test to compare *spo0A* site-directed mutants to the wild-type control (MC848) using GraphPad Prism v8.0.

### Western blotting

*C. difficile* strains were grown in BHIS supplemented with 5 µg ml^-1^ thiamphenicol, 0.2% fructose, and 0.1% taurocholate. Cultures were then diluted, grown to an OD_600_ of 0.5, and 250 µL of culture was plated on 70:30 agar. After 12 h, 5 ml of cells were scraped from agar, pelleted, and then washed with 1x PBS. Cells were resuspended in 1X sample buffer (10% glycerol, 5% 2-mercaptoethanol, 62.5 mM upper tris, 3% SDS, 5 mM PMSF) and lysed using a Biospec BeadBeater. Total protein concentration was then measured using a BCA protein assay kit (Pierce), and 4 µg of protein was separated by SDS-PAGE using pre-cast TGX 4-15% gradient gels (BioRad). Protein was transferred to a 0.45 µm nitrocellulose membrane, and Spo0A was detected using anti-Spo0A antibody [30]. Goat anti-mouse IgG Alexa fluor 488 (Invitrogen) was used as a secondary antibody, and western blots were visualized using a BioRad ChemiDoc MP System.

### Phos-tag blotting

*C. difficile* strains were cultured as described for western blotting. Cells from two plates for each strain were collected and pelleted. Cell pellets were suspended in 1 ml of 1X sample buffer (5% SDS, 93 mM Tris, 10% glycerol, 100 mM DTT). Protease Inhibitor Cocktail II (Sigma-Aldrich) was included in the sample buffer to inhibit protein degradation. Cells were lysed using a bead beater as described above. Total protein was measured using a BCA protein assay kit (Pierce). 10 μg protein aliquots were kept at 4°C or heated to 99°C for 10 min to dephosphorylate Spo0A prior to loading onto a 12.5% SuperSep Phos-tag gel (Fujifilm Wako)[50, 51]. Total protein was electrophoresed at 125 V for two hours at 4°C. The gel was rinsed three times in transfer buffer with 10% methanol and 10 mM EDTA to remove zinc present within the gel, and subsequently transferred to a low-fluorescence PVDF membrane (Thermo Scientific) in transfer buffer containing 10% methanol and 0.5% SDS overnight at 4°C. Western blot analysis was conducted with anti-FLAG M2 antibody (Sigma-Aldrich), followed by goat anti-mouse Alexa Fluor 488-conjugated antibody (Invitrogen) as the secondary. Imaging was performed using the BioRad ChemiDoc MP system.

### Two-hybrid luciferase assays

Two-hybrid assays were performed using a *C. difficile* codon-optimized split luciferase system previously described [52, 53]. *C. difficile* strains were grown in 70:30 broth supplemented with 2 µg ml^-1^ thiamphenicol. Cultures were grown to an OD_600_ of 0.8 – 0.9, then induced with 50 ng ml^-1^ anhydrous tetracycline for 1 hour. After induction, the OD_600_ were recorded, and 100 µL of each culture was added in technical duplicate to a chimney-style 96 well plate. Split-luciferase assay was then performed per manufacturer’s instructions (Promega). Luminescence output was immediately recorded at 135 nm using a BioTek plate reader. Output was normalized to cell density (OD_600_). A one-way ANOVA with Dunnett’s multiple comparisons test was performed to determine the statistical significance of luminescence outputs of the site-directed mutants relative to the wildtype using GraphPad Prism v8.0.

## ACKNOWLEDGEMENTS

We are appreciative for the gift of the anti-Spo0A antibody from Aimee Shen. We give special thanks to members of the McBride lab for suggestions during the completion of this work and preparation of this manuscript. This research was supported by the U.S. National Institutes of Health through research grants AI116933 and AI156052 to S.M.M. and GM008490 to M.A.D. The content of this manuscript is solely the responsibility of the authors and does not necessarily reflect the official views of the National Institutes of Health.

