## Supplemental Material for "Identification of functional Spo0A residues critical for sporulation in *Clostridioides difficile*"

**Supplemental Table 1.** Luminescence outputs from split-luciferase assay.

| **Strain** | **Average LU/$OD_{600}$** |
|---|---|
| Positive control (bitLuc$^{opt}$) | 9422.6 ± 1035.5 |
| Negative control (SmBit-LgBit) | 1051.6 ± 328.0 |
| Spo0A-SmBit | 1820.5 ± 692.9 |
| Spo0A-LgBit | 845.7 ± 188.2 |
| Spo0A-SmBit-Spo0A-LgBit | 934244.3 ± 47268.6 |
| Spo0A D10A-SmBit | 2517.3 ± 456.7 |
| Spo0A D10A-LgBit | 1307.8 ± 64.2 |
| Spo0A D10A-SmBit-Spo0A D10A-LgBit | 341759.3 ± 145113 |
| Spo0A D11A-SmBit | 969.4 ± 195.5 |
| Spo0A D11A-LgBit | 1378.3 ± 44.6 |
| Spo0A D11A-SmBit-Spo0A D11A-LgBit | 399696.3 ± 145900 |
| Spo0A D56A-SmBit | 998.9 ± 141.4 |
| Spo0A D56A-LgBit | 1203.4 ± 370.1 |
| Spo0A D56A-SmBit-Spo0A D56A-LgBit | 242346.6 ± 89320.3 |
| Spo0A I58A-SmBit | 2200.3 ± 788.8 |
| Spo0A I58A-LgBit | 1609.2 ± 199.9 |
| Spo0A I58A-SmBit-Spo0A I58A-LgBit | 442895.4 ± 269303 |
| Spo0A K108A-SmBit | 2648.6 ± 196.3 |
| Spo0A K108A-LgBit | 1592.4 ± 54.6 |
| Spo0A K108A-SmBit-Spo0A K108A-LgBit | 542192.7 ± 443215 |

**Supplemental Table 2**. Cloning and vector construction details

pMC566: A 1.2 kb *spo0A* PCR product amplified with primers oMC1249/oMC1250 was cloned into pMC123 using BamHI/EcoRI sites.

pMC567: A single amino acid mutation (D56A) within a 1.2 kb *spo0A* PCR product amplified with primers oMC1249/oMC1250 was made in a SOEing PCR reaction with two fragments generated by using primer set oMC1251/oMC1252 and cloned into pMC123 using BamHI/EcoRI sites.

pMC656: A single amino acid mutation (N12A) within a 1.2 kb *spo0A* PCR product amplified with primers oMC1249/oMC1250 was made in a SOEing PCR reaction with two fragments generated by using primer set oMC1513/oMC1514 and cloned into pMC123 using BamHI/EcoRI sites.

pMC657: A single amino acid mutation (K13A) within a 1.2 kb *spo0A* PCR product amplified with primers oMC1249/oMC1250 was made in a SOEing PCR reaction with two fragments generated by using primer set oMC1515/oMC1516 and cloned into pMC123 using BamHI/EcoRI sites.

pMC663: A single amino acid mutation (I58A) within a 1.2 kb *spo0A* PCR product amplified with primers oMC1249/oMC1250 was made in a SOEing reaction with two fragments generated by using primer set oMC1519/oMC1520 and cloned into pMC123 using BamHI/EcoRI sites.

pMC674: A 1.3 kb *spo0A* PCR product with C-terminal 3xFLAG amplified with primers oMC1249/oMC1547 was cloned into pMC123 using BamHI/EcoRI sites.

pMC684: A single amino acid mutation (V18A) within a 1.2 kb *spo0A* PCR product amplified with primers oMC1249/oMC1250 was made in a SOEing PCR reaction with two fragments generated by using primer set oMC1517/oMC1518 and cloned into pMC123 using BamHI/EcoRI sites.

pMC685: A single amino acid mutation (H61A) within a 1.2 kb *spo0A* PCR product amplified with primers oMC1249/oMC1250 was made in a SOEing PCR reaction with two fragments generated by using primer set oMC1583/oMC1584 and cloned into pMC123 using BamHI/EcoRI sites.

pMC697: A 1.2 kb *spo0A* C16A allele (TGT -> GCT) synthesized by Genscript was cloned into pMC123 using BamHI/EcoRI sites.

pMC698: A 1.2 kb *spo0A* E21A allele (GAG -> GCT) synthesized by Genscript was cloned into pMC123 using BamHI/EcoRI sites.

pMC699: A 1.2 kb *spo0A* A35S allele (GCT -> TCT) synthesized by Genscript was cloned into pMC123 using BamHI/EcoRI sites.

pMC700: A 1.2 kb *spo0A* P60A allele (CCA -> GCA) synthesized by Genscript was cloned into pMC123 using BamHI/EcoRI sites.

pMC701: A 1.2 kb *spo0A* A87S allele (GCA -> TCA) synthesized by Genscript was cloned into pMC123 using BamHI/EcoRI sites.

pMC702: A 1.2 kb *spo0A* V88A allele (GTA -> GCA) synthesized by Genscript was cloned into pMC123 using BamHI/EcoRI sites.

pMC703: A 1.2 kb *spo0A* G89A allele (GGT -> GCT) synthesized by Genscript was cloned into pMC123 using BamHI/EcoRI sites.

pMC704: A 1.2 kb *spo0A* K108A allele (AAG -> GCA) synthesized by Genscript was cloned into pMC123 using BamHI/EcoRI sites.

pMC742: A 1.2 kb *spo0A* D91A allele (GAT -> GCT) synthesized by Genscript was cloned into pMC123 using BamHI/EcoRI sites.

pMC768: A 1.2 kb *spo0A* M59A allele (ATG -> GCA) synthesized by Genscript was cloned into pMC123 using BamHI/EcoRI sites.

pMC769: A 1.2 kb *spo0A* L62A allele (CTA -> GCA) synthesized by Genscript was cloned into pMC123 using BamHI/EcoRI sites.

pMC770: A 1.2 kb *spo0A* K92A allele (AAG -> GCA) synthesized by Genscript was cloned into pMC123 using BamHI/EcoRI sites.

pMC771: A 1.2 kb *spo0A* P109A allele (CCA -> GCA) synthesized by Genscript was cloned into pMC123 using BamHI/EcoRI sites.

pMC922: Two PCR fragments were generated with oMC2437/2439 (950 bp) + oMC2441/2442 (600 bp) to create *spo0A*-SmBit-LgBit and Gibson assembled as BamHI/SacI into pAP118.

pMC924: Three PCR fragments were generated with oMC2443/2356 (150 bp) + oMC2354/2447 (950 bp) + oMC2445/2442 (550 bp) to create SmBit-*spo0A*-LgBit and Gibson assembled as BamHI/SacI into pAP118.

pMC930: Two PCR fragments were combined in SOEing PCR reaction from oMC2443/2449 and oMC2442/2448 (600 bp) and Gibson assembled into pAF256 to create SmBit-LgBit fusion.

pMC932: Two PCR fragments were combined in SOEing PCR reaction from oMC2437/2439 and oMC2441/2356 (1050 bp) and cloned into the SacI/PvuI sites of pAF257 to create *spo0A*-SmBit.

pMC944: Two PCR fragments were combined in SOEing PCR reaction from oMC2354/2447 and oMC2445/2442 (1850 bp) and cloned into the BamHI/PvuI sites of pMC932 to create *spo0A*-SmBit-*spo0A*-LgBit.

pMC965: A 1.2 kb *spo0A* D11A allele (GAC -> GCA) synthesized by Genscript was cloned into pMC123 using BamHI/EcoRI sites.

pMC966: A 1.2 kb *spo0A* D14A allele (GAT -> GCT) synthesized by Genscript was cloned into pMC123 using BamHI/EcoRI sites.

pMC967: A 1.2 kb *spo0A* F15A allele (TTT -> GCT) synthesized by Genscript was cloned into pMC123 using BamHI/EcoRI sites.

pMC968: A 1.2 kb *spo0A* Q17A allele (CAG -> GCT) synthesized by Genscript was cloned into pMC123 using BamHI/EcoRI sites.

pMC969: A 1.2 kb *spo0A* L19A allele (TTA -> GCA) synthesized by Genscript was cloned into pMC123 using BamHI/EcoRI sites.

pMC970: A 1.2 kb *spo0A* D111A allele (GAT -> GCT) synthesized by Genscript was cloned into pMC123 using BamHI/EcoRI sites.

pMC971: *spo0A* D56A-3XFLAG (SOEing product from oMC1249/1252 and oMC1251/1547) was made and cloned into pMC123 using BamHI/EcoRI.

pMC975: A 1.2 kb *spo0A* K36A allele (AAG -> GCA) synthesized by Genscript was cloned into pMC123 using BamHI/EcoRI sites.

pMC976: A 1.2 kb *spo0A* Q90A allele (CAA -> GCA) synthesized by Genscript was cloned into pMC123 using BamHI/EcoRI sites.

pMC986: A 1.2 kb *spo0A* F110A allele (TTT -> GCT) synthesized by Genscript was cloned into pMC123 using BamHI/EcoRI sites.

pMC1055: A 1.2 kb *spo0A* S86A allele (TCA -> GCA) synthesized by Genscript was cloned into pMC123 using BamHI/EcoRI sites.

pMC1088: A 1.2 kb *spo0A* D10A allele (GAT -> GCT) synthesized by Genscript was cloned into pMC123 using BamHI/EcoRI sites.

pMC1097: An 850 bp *spo0A* D10A PCR fragment was amplified from pMC1088 using oMC2437/oMC2439 and was Gibson assembled into pAF256.

pMC1098: An 850 bp *spo0A* D11A PCR fragment was amplified from pMC965 using oMC2437/oMC2439 and was Gibson assembled into pAF256.

pMC1099: An 850 bp *spo0A* D56A PCR fragment was amplified from pMC567 using oMC2437/oMC2439 and was Gibson assembled into pAF256.

pMC2000: An 850 bp *spo0A* I58A PCR fragment was amplified from pMC663 using oMC2437/oMC2439 and was Gibson assembled into pAF256.

pMC2001: An 850 bp *spo0A* K108A PCR fragment was amplified from pMC704 using oMC2437/oMC2439 and was Gibson assembled into pAF256.

pMC2002: An 850 bp *spo0A* D10A PCR fragment was amplified from pMC1088 using oMC2354/oMC2447 and was Gibson assembled into pAF257.

pMC2003: An 850 bp *spo0A* D11A PCR fragment was amplified from pMC965 using oMC2354/oMC2447 and was Gibson assembled into pAF257.

pMC2004: An 850 bp *spo0A* D56A PCR fragment was amplified from pMC567 using oMC2354/oMC2447 and was Gibson assembled into pAF257.

pMC2005: An 850 bp *spo0A* I58A PCR fragment was amplified from pMC663 using oMC2354/oMC2447 and was Gibson assembled into pAF257.

pMC2006: An 850 bp *spo0A* K108A PCR fragment was amplified from pMC704 using oMC2354/oMC2447 and was Gibson assembled into pAF257.

pMC2007: An 850 bp *spo0A* D10A PCR fragment was amplified from pMC1088 using oMC2437/oMC2439 and was Gibson assembled into pAP118. An 850 bp *spo0A* D10A PCR fragment was then amplified from pMC1088 using oMC2354/oMC2447 and cloned into pAP118 using NotI and PvuI sites.

pMC2008: An 850 bp *spo0A* D11A PCR fragment was amplified from pMC965 using oMC2437/oMC2439 and was Gibson assembled into pAP118. An 850 bp *spo0A* D11A PCR fragment was then amplified from pMC965 using oMC2354/oMC2447 and cloned into pAP118 using NotI and PvuI sites.

pMC2009: An 850 bp *spo0A* D56A PCR fragment was amplified from pMC567 using oMC2437/oMC2439 and was Gibson assembled into pAP118. An 850 bp *spo0A* D56A PCR fragment was then amplified from pMC567 using oMC2354/oMC2447 and cloned into pAP118 using NotI and PvuI sites.

pMC2010: An 850 bp *spo0A* I58A PCR fragment was amplified from pMC663 using oMC2437/oMC2439 and was Gibson assembled into pAP118. An 850 bp *spo0A* I58A PCR fragment was then amplified from pMC663 using oMC2354/oMC2447 and cloned into pAP118 using NotI and PvuI sites.

pMC2011: An 850 bp *spo0A* K108A PCR fragment was amplified from pMC704 using oMC2437/oMC2439 and was Gibson assembled into pAP118. An 850 bp *spo0A* K108A PCR fragment was then amplified from pMC704 using oMC2354/oMC2447 and cloned into pAP118 using NotI and PvuI sites.

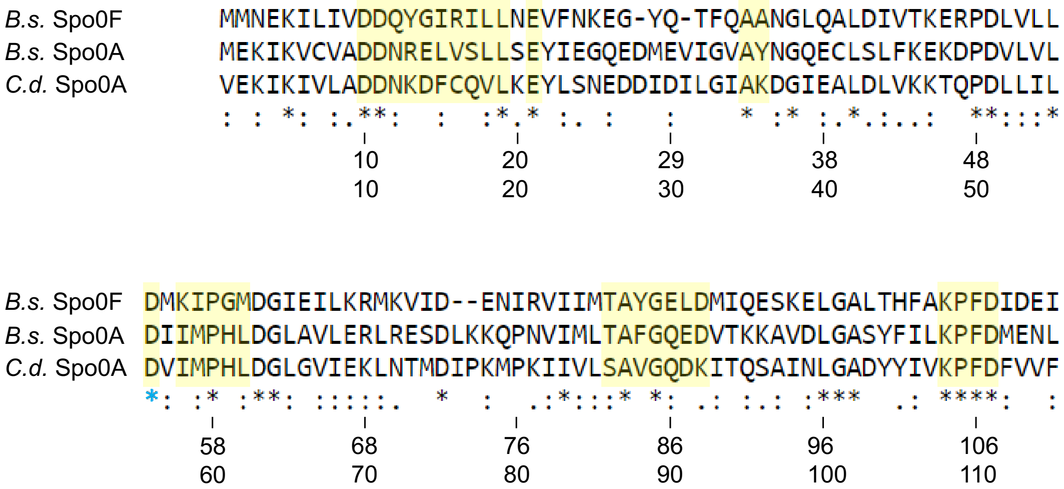


**Supplementary Figure 1. Alignment of receiver domain residues of *B. subtilis* Spo0A and Spo0F, and *C. difficile* Spo0A.** Spo0A receiver domain of *C. difficile* aligned to *B. subtilis* Spo0F and receiver domain of Spo0A using Clustal Omega (*B.s.* = *B. subtilis*; *C.d.* = *C. difficile*). Conserved residues chosen for mutation in *C. difficile* are highlighted in yellow. Amino acid sequences of Spo0F (BSU_37130) and of the Spo0A receiver domains for *B. subtilis str. 168* (BSU_24220, top), and *C. difficile* 630 (CD630_12140, bottom). The blue star (*) is the conserved site of phosphorylation.

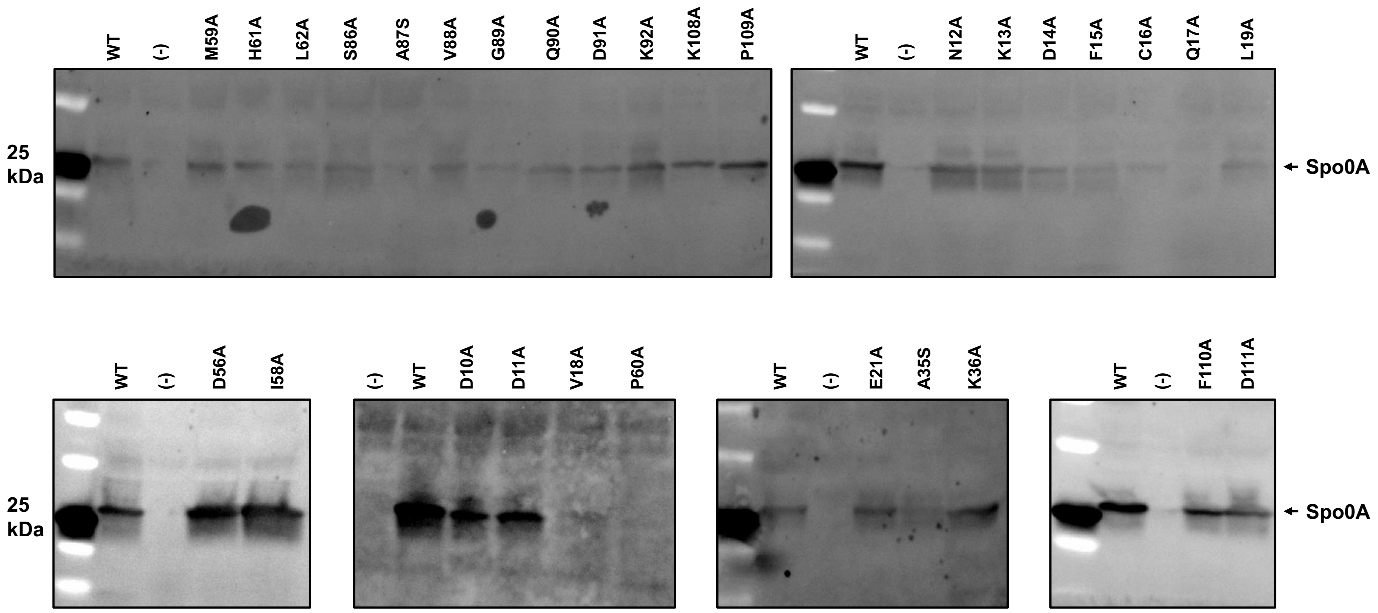


**Supplementary Figure 2. Stability of Spo0A mutant alleles.** SDS-PAGE western blot analysis of the 30 Spo0A point mutants to assess their stability using anti-Spo0A antibody. Strains harvested after 12 h growth on 70:30 sporulation medium and 4 $\mu$g of total protein was loaded for each sample. A wild type (WT) positive control, *spo0A*::*erm* pSpo0A (MC848) and the *spo0A*::*erm* pMC123 (MC855) negative control strain (-) are included for each western blot.

A)

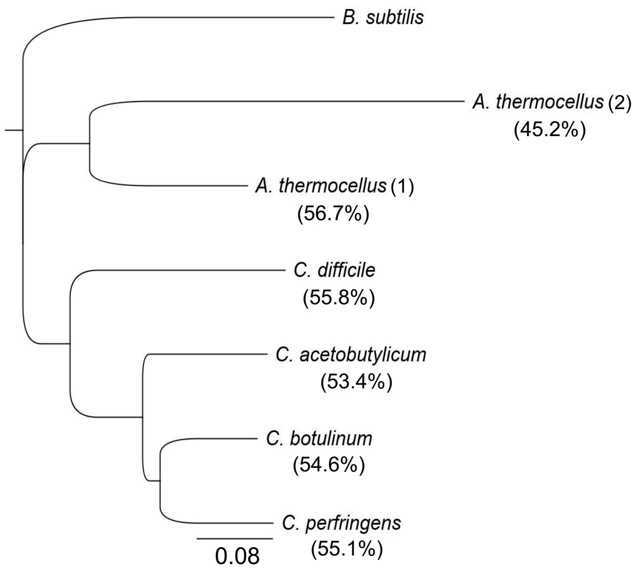


B) Spo0A % identity

| | ***A.t.* 2** | ***B.s.*** | ***A.t.* 1** | ***C.d.*** | ***C.p.*** | ***C.b.*** | ***C.a.*** |
|---|---|---|---|---|---|---|---|
| ***A.t.* 2** | **-** | **45.2** | **54.9** | **47.6** | **49.8** | **47.8** | **48.2** |
| ***B.s.*** | **45.2** | **-** | **56.7** | **55.4** | **55.1** | **54.6** | **53.4** |
| ***A.t.* 1** | **54.9** | **56.7** | **-** | **56.7** | **61.2** | **62.5** | **59.1** |
| ***C.d.*** | **47.8** | **55.8** | **57.1** | **-** | **60.2** | **64.6** | **61.3** |
| ***C.p.*** | **49.8** | **55.1** | **61.2** | **59.8** | **-** | **78.6** | **75.1** |
| ***C.b.*** | **47.8** | **54.6** | **62.5** | **64.1** | **78.6** | **-** | **76.5** |
| ***C.a.*** | **48.2** | **53.4** | **59.1** | **60.9** | **75.1** | **76.5** | **-** |

**Supplementary Figure 3. Spo0A divergence in Firmicutes. A)** Dendrogram of full-length Spo0A protein coding regions rooted to the outgroup *B. subtilis* Spo0A. Percentage identity of each species' Spo0A protein sequence is shown relative to *B. subtilis*. Spo0A alignment and dendrogram tree constructed using MUSCLE Alignment plugin and Geneious Tree Builder in Geneious Prime 2020.2.2. **B)** Heatmap of the comparisons of percent identities of Spo0A from each species in **(A)**. *B.s.* = *B. subtilis*, *A.t.*1 = *A. thermocellus* Spo0A 1, *A.t.* 2 = *A. thermocellum* Spo0A 2, *C.d.* = *C. difficile*, *C.p.* = *C. perfringens*, *C.b.* = *C. botulinum*, and *C.a.* = *C. acetobutylicum*.
